# Reversing Pathophysiology in Fragile X Syndrome Mice by Promoting PGC-1α and Mitochondrial Functions

**DOI:** 10.64898/2026.04.13.718178

**Authors:** Anirudh Acharya, Vipendra Kumar, Kwan Young Lee, Matthew S Babik, Gaurav Goswami, Kingsley A. Boateng, Austin J. Cyphersmith, Justin S Rhodes, Nien-Pei Tsai

## Abstract

Fragile X syndrome (FXS) is the leading cause of intellectual disabilities and autism, but a disease-modifying strategy remains unavailable. Recent studies have suggested reduced mitochondrial functions in FXS. However, the mechanisms underlying mitochondrial defects and their impact on FXS pathophysiology remain largely unclear. Here, we reveal a reduction in the mitochondrial master regulator peroxisome proliferator-activated receptor gamma coactivator 1-alpha (PGC-1α) in the mouse model of FXS, the *Fmr1* knockout (KO) mice. We show that this impairment is caused by the inactivity of the transcription factor cAMP-response element-binding protein (CREB) in *Fmr1* KO mice. Using the small molecule ZLN005, which induces AMP-activated protein kinase (AMPK)- and CREB-dependent elevation of PGC-1α in *Fmr1* KO mice, we observed significantly increased mitochondrial functions and dynamics in cultured neurons in vitro and in the hippocampus in vivo. Furthermore, ZLN005 elicited a wide range of beneficial effects in *Fmr1* KO mice, including enhanced inhibitory synaptic transmission, reduced circuit hyperexcitability, improved hippocampal synaptic plasticity, reduced cortical gamma-band oscillations, and improved interhemispheric coherence. Most importantly, we observed improved cognition and reduced autism-like behaviors in ZLN005-treated *Fmr1* KO mice. Together, our findings identify AMPK-CREB signaling and PGC-1α as promising and selective therapeutic targets for FXS and reveal the broad impact of restoring PGC-1α on FXS pathophysiology.

**One Sentence Summary:** Promoting PGC-1α Reverses FXS Pathophysiology.

## INTRODUCTION

Fragile X syndrome (FXS) is the most common genetic cause of autism spectrum disorder (ASD), arising from silencing of the *Fmr1* gene on the X chromosome, which encodes the fragile X messenger ribonucleoprotein (FMRP). FMRP is a translational repressor that regulates the expression of numerous neuronal proteins. Its absence disrupts various aspects of brain development and function, including synapse formation and maturation, neuronal excitability, plasticity, sensory processing, and cognition (*1, 2*). As a result, FXS patients exhibit intellectual disabilities, and the condition predominantly affects males due to a single copy of the *Fmr1* gene. Given FMRP’s complex role in multiple neuronal signaling pathways, no single drug has been shown to address the wide range of FXS symptoms. Additionally, experimental treatments often cause undesirable side effects. Currently, no medications are clinically approved to treat FXS, motivating our research to identify novel pharmacological targets that may improve patients’ quality of life.

Mitochondrial dysfunction is common in neurological disorders (*3–6*). Neurons, unlike most cell types, have evolved to regulate mitochondrial functions dynamically, enabling rapid mitochondrial plasticity in response to activity. Precise mitochondrial localization in axons and dendrites is essential for energy homeostasis and neuronal function (*7*). In FXS, mitochondrial abnormalities have been reported (*8–11*). Because mitochondria are crucial for calcium buffering, these abnormalities can also disrupt molecular signaling pathways in FXS. Interventions aimed at improving mitochondrial function have shown therapeutic potential in FXS (*12*). However, the specific targets and broad impact on FXS pathology remain unclear.

Peroxisome proliferator-activated receptor-γ coactivator-1α (PGC-1α) is a master regulator of mitochondrial functions (*13*). As a transcriptional co-activator, PGC-1α is also essential for the expression of various genes that support neuronal functions (*14*). Previous studies have shown that PGC-1α regulates genes involved in mitochondrial fusion, such as *Mfn1* and *Mfn2* (*15*), and promotes enzymes such as superoxide dismutase that decrease the levels of reactive oxygen species (*16*). In the brain, reduced PGC-1α expression drastically decreases calcium-binding proteins, such as parvalbumin (PV) (*17*) and synaptotagmin (*18*), and alters neuronal physiology, including firing properties and plasticity (*19, 20*). There are indications that changes in expression of calcium buffering proteins could trigger alterations in mitochondrial physiology, leading to autism (*21*). These prior studies suggest the critical role of PGC-1α in regulating neuronal physiology through mitochondria. However, the relationship between PGC-1α and FXS remains elusive.

In this study, we revealed a reduction of PGC-1α in the mouse model of FXS, the *Fmr1* KO mice. We further uncovered that this reduction is caused by the inactivity of transcription factor cAMP-response element-binding protein (CREB) in *Fmr1* KO mice. Using a small molecule ZLN005 that promotes phosphorylation of AMP-activated protein kinase (AMPK), AMPK-dependent activation of CREB and CREB-dependent PGC-1α expression in *Fmr1* KO mice, we observed drastic changes of mitochondrial functions in cultured *Fmr1* KO neurons in vitro and the mitochondrial dynamics in hippocampus of *Fmr1* KO mice in vivo. In addition, we also observed many positive impacts on *Fmr1* KO mice, including enhanced inhibitory synaptic transmission, reduced brain hyperexcitability, improved hippocampal synaptic plasticity, reduced cortical LFP high-gamma band power, enhanced interhemispheric coherence in the theta band, improved cognition and reduced autism-like behavior in *Fmr1* KO mice. Importantly, we observed no significant effects in WT mice after ZLN005 treatment, likely due to basally active AMPK-CREB signaling in WT neurons. In summary, our study identifies AMPK-CREB signaling and PGC-1α as novel and selective therapeutic targets for FXS and comprehensively illustrates the impact of restoring PGC-1α on FXS pathophysiology.

## RESULTS

### ZLN005 facilitates AMPK- and CREB-dependent expression of PGC-1α in *Fmr1* KO neurons

To begin studying the role of PGC-1α in FXS, we first compared the expression level of PGC-1α in WT and *Fmr1* KO cortical neuron cultures. As shown in Figure 1A, we observed a significant reduction of PGC-1α in *Fmr1* KO cultures. To reverse this downregulation, we employed ZLN005, a known PGC-1α activator, in both WT and *Fmr1* KO cortical neuron cultures. As shown in Figure 1B, ZLN005 (5 μM, 24 hrs) significantly elevates PGC-1α in *Fmr1* KO, but not WT, cultures. To understand the mechanism underlying selective elevation of PGC-1α in *Fmr1* KO, we study CREB. Previous studies have shown that activation of CREB can promote PGC-1α expression (*22*). To test whether ZLN005 modulates CREB activity and whether the effect is different between WT and *Fmr1* KO neurons, we measured CREB phosphorylation in WT and *Fmr1* KO cortical neuron cultures treated with ZLN005 (5 μM, 1 hr). As shown in Figure 1C, CREB phosphorylation is basally reduced in *Fmr1* KO neurons but can be selectively elevated by ZLN005. No significant effects were observed in WT neurons. To confirm that CREB is responsible for ZLN005-induced PGC-1α expression in *Fmr1* KO neurons, we administered ZLN005 (5 μM) and a CREB inhibitor 666-15 (20 nM) in *Fmr1* KO cortical neuron cultures for 24 hrs. As shown in Figure 1D, we found that 666-15 inhibits ZLN005-induced elevation of PGC-1α expression in *Fmr1* KO neurons. Together, these data suggest that ZLN005 selectively activates CREB to promote PGC-1α expression in *Fmr1* KO neurons.

**Figure 1.**
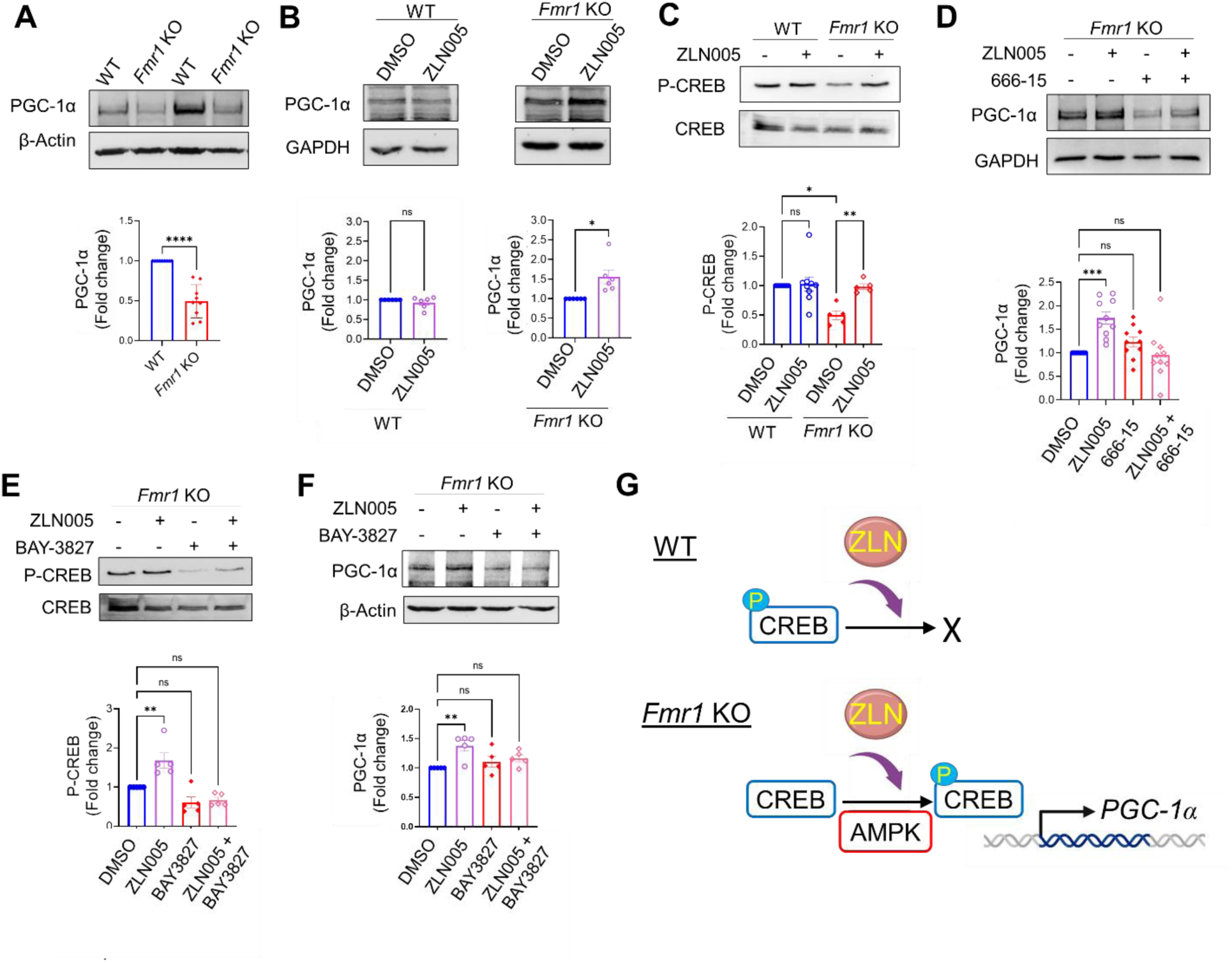
AMPK and CREB guide the expression of PGC-1α, and this mechanism is impaired in *Fmr1* KO mice. (**A**) Representative western blots of PGC-1α and β-actin from WT or *Fmr1* KO primary cortical neurons. The quantification of PGC-1α over β-actin was at the bottom. (n = 9 independent cultures). (**B**) Representative western blots of PGC-1α and GAPDH from WT or *Fmr1* KO primary cortical neurons treated with DMSO or ZLN005 (5 μM, 24 hrs). The quantification of PGC-1α over GAPDH was at the bottom. (n = 6 independent experiments). (**C**) Representative western blots of CREB and p-CREB from WT or *Fmr1* KO primary cortical neurons treated with DMSO or ZLN005 (5 μM, 1 hr). The quantification of p-CREB over CREB was at the bottom. (n = 5-9 independent experiments). (**D**) Representative western blots of PGC-1α and GAPDH from *Fmr1* KO primary cortical neurons treated with ZLN005 (5 μM) with or without 666-15 (20 nM) for 24 hrs. The quantification of PGC-1α over GAPDH was at the bottom. (n = 10 independent experiments). (**E**) Representative western blots of CREB and p-CREB from *Fmr1* KO primary cortical neurons treated with ZLN005 (5 μM, 24 hrs) with or without BAY-3827 (1 μM) during the last one hour of ZLN005 treatment. The quantification of p-CREB over CREB was at the bottom. (n = 5 independent experiments). (**F**) Representative western blots of PGC-1α and β-actin from *Fmr1* KO primary cortical neurons treated with ZLN005 (5 μM) with or without BAY-3827 (1 μM) for 24 hrs. The quantification of PGC-1α over GAPDH was at the bottom. (n = 10 independent experiments). (**G**) Summary diagram showing the selective effect of ZLN005 on *Fmr1* KO neurons to activate AMPK-CREB signaling-dependent elevation of PGC-1α. For data analysis, significance was determined by Student’s *t*-test (A, B) or two-way ANOVA with Tukey test (C-F). Data are represented as mean ± SEM with *P<0.05, **P<0.01, ***P<0.001, ns: non-significant.

We next asked how ZLN005 promotes CREB phosphorylation. Previous studies have suggested that ZLN005 functions in part through activating AMPK (*23*) and AMPK can phosphorylate CREB (*24*). To study AMPK, we treated *Fmr1* KO cortical neuron cultures with an AMPK inhibitor, BAY-3827 (1 μM, 1hr). As shown in Figure 1E, BAY-3827 inhibits the elevation of CREB phosphorylation in *Fmr1* KO neurons. As expected, treatment of BAY-3827 (1 μM) also inhibits the elevation of PGC-1α induced by ZLN005 (5 μM, 24 hrs) (Figure 1F). These findings suggest that ZLN005 activates AMPK-CREB signaling in *Fmr1* KO neurons to elevate PGC-1α expression, and the lack of effect in WT neurons is likely caused by basally elevated CREB phosphorylation (Figure 1G).

### ZLN005 promotes mitochondrial functions and dynamics in *Fmr1* KO mice

Mitochondrial defects are documented in FXS animal models and patient-derived cells (*8–11*). Because PGC-1α is a master mitochondrial regulator, we ask whether ZLN005 can improve mitochondrial functions in *Fmr1* KO mice. We first employed MitoTracker dye to determine mitochondrial membrane potential. As shown in Figure 2A, *Fmr1* KO neurons exhibit a lower uptake of MitoTracker when compared to WT neurons. Importantly, ZLN005 significantly improved the uptake of MitoTracker in *Fmr1* KO neurons without obvious effects in WT neurons. These results suggest that ZLN005 elevates mitochondrial membrane potential in *Fmr1* KO neurons. These data were supported by a ZLN005-induced elevation of multiple mitochondrial marker proteins in *Fmr1* KO neurons, including Voltage-dependent anion channels (VDAC), Mitofusin 2 (MFN2), Nuclear respiratory factor 1 (NRF1), Dynamin-Related Protein 1 (DRP1), and Cytochrome-c oxidase IV isoform (COX-IV) (Figure 2B). Altogether, these results suggest that ZLN005 can promote mitochondrial functions in *Fmr1* KO neurons.

**Figure 2.**
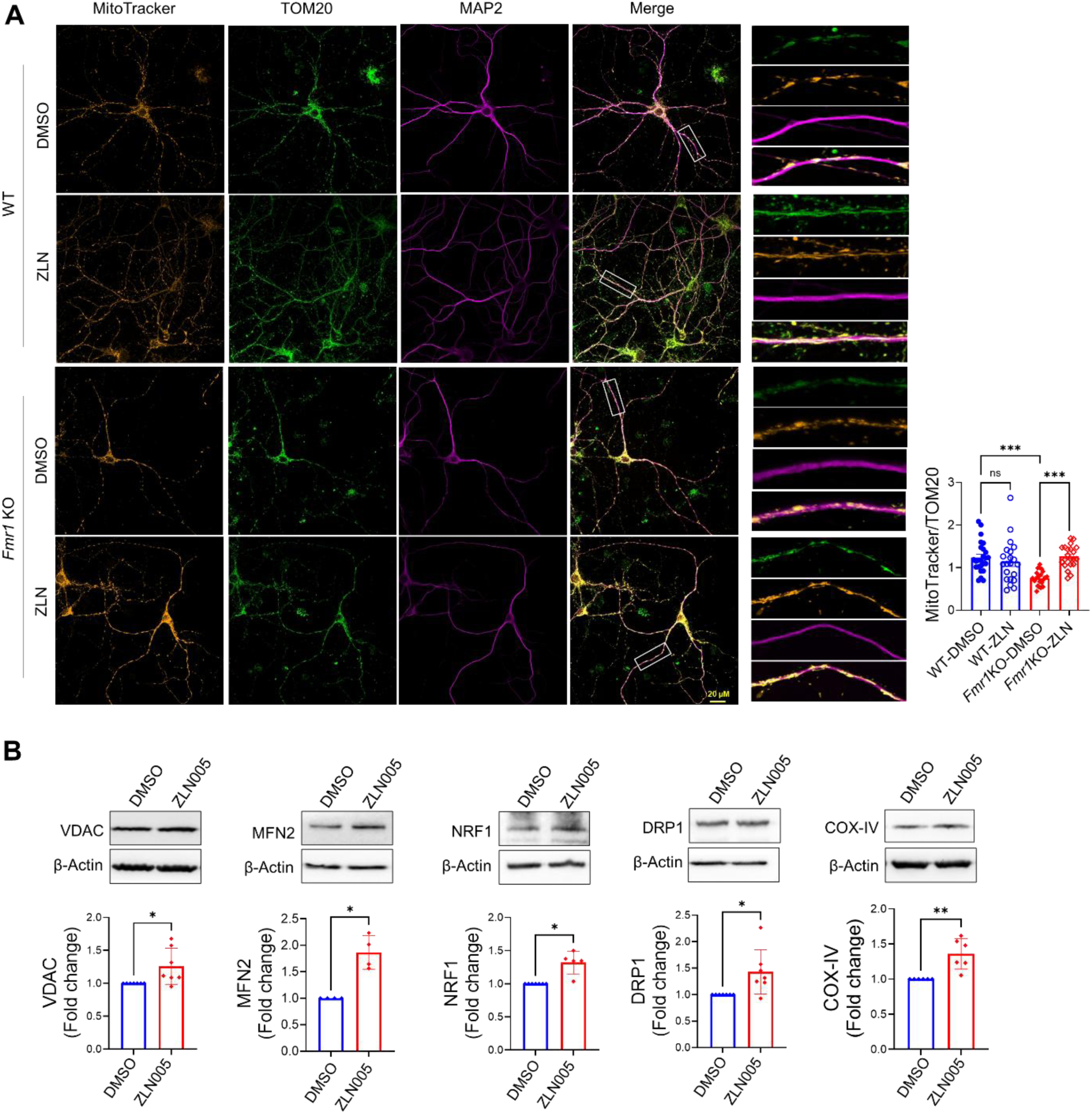
Promoting PGC-1α via ZLN005 improves mitochondrial properties in *Fmr1* KO mice. (**A**) Representative immunocytochemistry images of MitoTracker dye, mitochondrial marker TOM20, and dendritic marker MAP2 from WT or *Fmr1* KO cortical neurons treated with DMSO or ZLN005 (5 μM, 24 hrs). Selective, enlarged dendritic areas were shown on the right. Quantification of MitoTracker over TOM20 was also shown on the right (n = 21-25 cells per condition). Scale bar: 20 µm. (**B**) Representative western blots of VDAC, MFN2, NRF1, DRP1, COX-IV and β-actin from *Fmr1* KO primary cortical neurons treated with DMSO or ZLN005 (5 μM, 24 hrs). The quantification of each protein over β-actin was shown at the bottom. (n = 4-7 independent experiments). Significance was determined by Student’s *t*-test. Data are represented as mean ± SEM with *P<0.05, **P<0.01, ns: non-significant.

To test the effect of ZLN005 in mice in vivo, we next performed serial block-face scanning electron microscopy (SBF-SEM) on WT and *Fmr1* KO mice at 7 weeks of age after daily treatments with saline or ZLN005 (10 mg/kg) for 7 days. Figure 3A shows a schematic representation of ZLN005 treatment and the SBF-SEM image processing that focuses on hippocampal CA1 pyramidal neurons. Figure 3B shows our image annotation and segmentation pipeline. This pipeline was achieved using our custom, in-house-developed Attention-based UNET machine learning model that segments mitochondria and nuclei in our SBF-SEM data (Figure S1A). To assess our model’s ability to accurately identify their corresponding organelles, we quantified the precision, recall, F1 score, and Dice coefficients for both mitochondrial and nuclear models (Figure S1B). Loss function curves for mitochondrial and nuclear model training were generated, showing a steep decline with increased training epochs as the models learned primary features (Figure S1B). We generated new images that the model had not been trained on and used this dataset to assess our performance metrics (Figure S1B). We deemed these metrics sufficient relative to the level of error in manual annotations. Although the performance metrics for the nuclei segmentation model were lower than those for the mitochondrial model, it was still sufficient to segment images successfully (Figure S1D). The model encountered some difficulty classifying patches from the center of the nuclei. To overcome this limitation, we employed patch-based inference on the full SBF-SEM images post-training. This strategy resolved the issue by leveraging a larger sampling space than the training data alone provides. Following the successful testing of our model, the cell bodies in the hippocampal subfield images were segmented semi-automatically using the Microscopy Image Browser software (*25*). The segmented binary masks of mitochondria, nuclei, and the cell body model were exported to Amira software for 3D reconstruction and quantitative analysis. Figure 3C shows a 3D-reconstructed WT hippocampal CA1 neuron with a cell body, nucleus, and mitochondria. An example video showing the 3D mitochondrial reconstruction in the cell body along the z-axis was provided in Movie S1.

**Figure 3.**
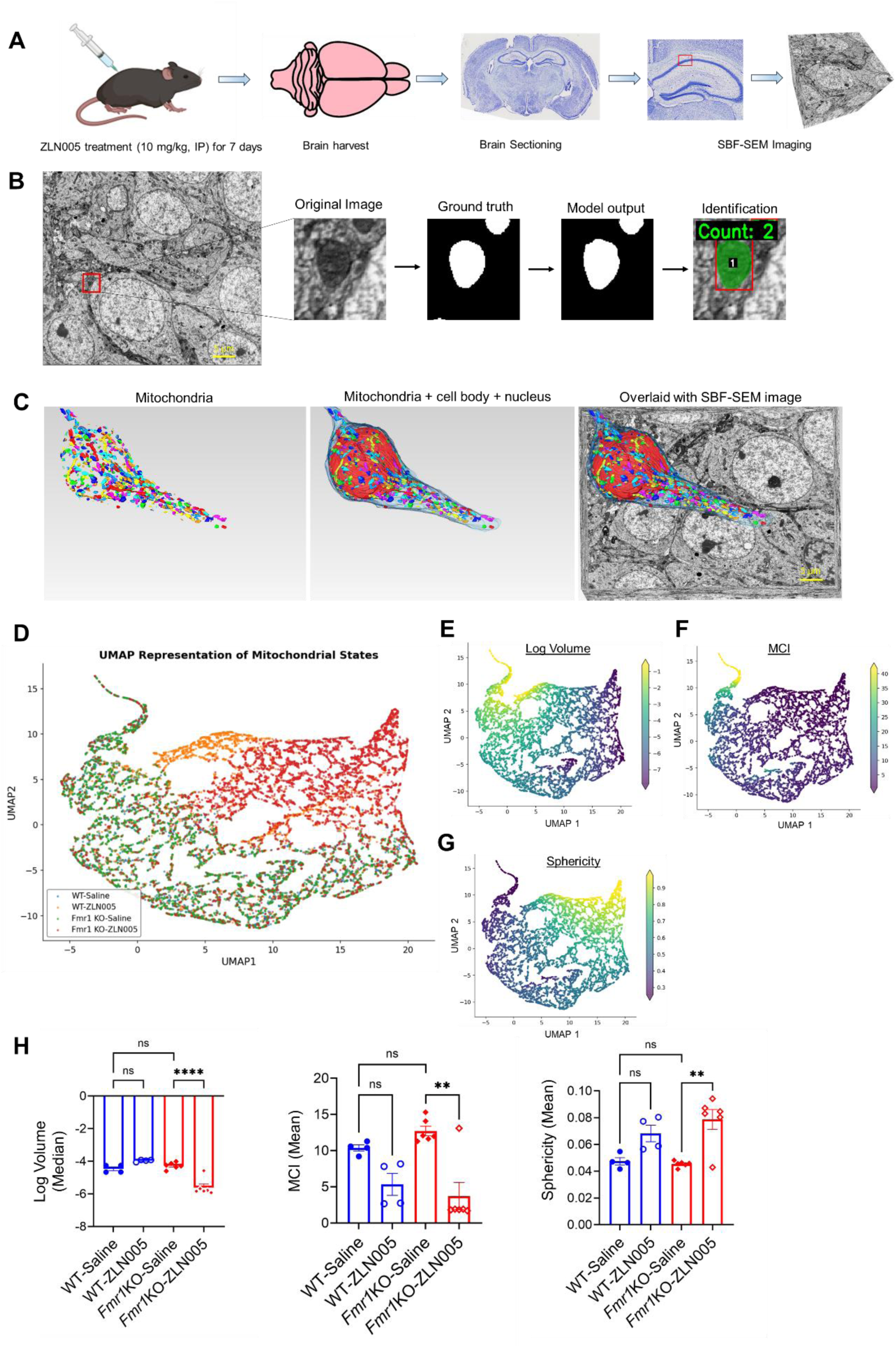
**ZNL005 induces genotype-specific remodeling of mitochondrial morphology in *Fmr1* KO neurons, revealed by attention U-Net-based volumetric SBF-SEM segmentation.** (**A**) The experimental design illustrating SBF-SEM procedures. WT and *Fmr1* KO mice were administered daily saline or ZLN005 (10 mg/kg, i.p.) for 7 days. After treatment, the mice were transcardially perfused and rapidly fixed and the brains were harvested. Two hundred µm-thick coronal sections were prepared, from which hippocampi were dissected and processed for SBF-SEM imaging. The pyramidal cell layer of each hippocampus was later imaged by SBF-SEM. (**B**) Representative SBF-SEM micrograph of hippocampal pyramidal layer (left). The red box on the micrograph shows a region of interest, a 200×200 pixel patch used for training the attention U-Net model. The segmentation pipeline was on the right, including original SBF-SEM image, the binary ground truth showing annotated mitochondria used for model training, the output generated by the model, and an image overlay after mitochondria identification. (**C**) Representative 3D reconstruction showing individual segmented mitochondria from the cell body of a WT CA1 neuron in the volumetric SBF-SEM images (left). Mitochondria and nucleus reconstructed within a neuronal cell body (middle) and the overlay of 3D reconstructed mitochondria on the original SBF-SM image (right). Scale bar: 5 µm. (**D**) UMAP embedding of mitochondria from all four experimental groups (color coded) with three morphometric features: log-transformed volume, MCI and sphericity into a group-colored plot. (**E-G**) Same embedding of all the mitochondria with individual morphometirc features. (**H**) Quantifications of log-transformed mitochondrial volume, MCI and sphericity across all four treatment groups. (n=4 and 6 for WT and *Fmr1* KO, respectively). Two-way ANOVA was used to calculate statistical significance. Data are presented as mean± SEM. **p < 0.01, ****p < 0.0001, ns: non-significant.

Previous studies have revealed that mitochondrial shapes vary depending on their sub-cellular localization in the neurons (*26*). Therefore, we calculated the log-transformed volume, mitochondrial complexity index (MCI), and sphericity to account for their complex and highly variable morphology. Next, to understand the relationships among these morphometric features across genotypes and treatments, we embedded all three features (log-transformed volume, MCI, and sphericity) into a multi-dimensional space using Uniform Manifold Approximation and Projection (UMAP), where each point indicates individual mitochondria. The colored UMAP embedding shows that mitochondria from all four experimental groups occupy overlapping yet regionally differentiated zones, organized along a size gradient from the upper left (large) to the lower right (small) and a shape gradient from the lower left (elongated) to the upper right (round) (Figure 3D). Additionally, to improve visualization, we created UMAP plots of individual morphometric features across all four treatment groups, showing heat maps of log-transformed volume, MCI, and sphericity (Figures 3E-3G). The manifold reveals that WT-saline (blue) and *Fmr1* KO-saline (green) mitochondria showed substantial spatial overlap in the lower-left part of the embedding. Interestingly, a morphologically extreme subpopulation was present as an elongated ribbon-like distribution in the upper-left region, populated mainly by the *Fmr1* KO-saline group, and characterized as highly complex, large, elongated mitochondria. Mitochondria from the WT-ZLN005 treated group (orange) substantially occupied the upper-central region in the UMAP plot (large, elongated mitochondria), and had less overlap with other groups, suggesting that ZLN005 treatment caused a distinctive shift in mitochondrial state in WT mice. On the other hand, mitochondria from ZLN005-treated *Fmr1* KO mice (red) were predominantly distributed in the lower right region of the plot (small, round mitochondria) and failed to colocalize with the WT-ZLN005 cluster. Overlaying the log-transformed mitochondrial volume on the UMAP embedding revealed a continuous size gradient from the upper left to the lower right of the manifold (Figures 3D and 3E). Although there was no statistically significant difference between WT-saline and *Fmr1* KO-saline, or between WT-saline and WT-ZLN005, ZLN005 treatment in *Fmr1* KO showed a significant reduction in median log volume (Figure 3F). The MCI overlay on the same UMAP embedding exhibited high MCI values (>25) for the upper-left ribbon subpopulation predominantly occupied by *Fmr1* KO-saline mitochondria, and the remaining mitochondria from all four groups had uniformly low MCI (<10) (Figures 3D and 3F). When cross-referenced with the UMAP of MCI with group-colored panels, it was revealed that ZLN005 treatment significantly reduced mean MCI in *Fmr1* KO neurons (p<0.01), whereas WT neurons did not show any significant difference (Figure 3H). Overlaying sphericity to colored UMAP showed a left-to-right shift (elongated ≈ 0.25–0.35 to round ≈ 0.7–0.9) in *Fmr1* KO-ZLN005 neurons (Figure 3D and 3G). The ZLN005-treated *Fmr1* KO neurons showed significantly increased sphericity as compared to the *Fmr1* KO saline-treated group (Figure 3H), but there was no significant effect of ZLN005 treatment on WT. Reduced mitochondrial volume, decreased MCI, and increased sphericity all indicate quality-control fission, higher metabolic rate and early-stage mitochondrial biogenesis (*27, 28*). Because smaller mitochondria also support neurotransmission and synaptic plasticity (*29*), our data suggest that ZLN005 induces *Fmr1* KO-specific elevation in mitochondrial functions that has the potential to strengthen brain circuit connectivity and maturation.

### ZLN005 ameliorates hyperexcitability phenotypes in *Fmr1* KO mice

FXS patients and the *Fmr1* KO mice both show abnormal up-regulation of excitability (*30–32*) with seizure diagnosed in 15-40% of the patients (*33, 34*) and hyperexcitability-associated behavioral symptoms observed in over 90% of the patients (*35*). To determine whether ZLN005 can reduce hyperexcitability phenotypes in *Fmr1* KO mice, we injected the mice with kainic acid (30 mg/kg i.p.) to chemically induce seizures (*36*) following daily treatments with saline or ZLN005 (2.5 mg/kg) for 7 days. After the kainic acid injections, the mice were closely observed in real time for one hour and the intensity of seizures was assessed by a modified Racine’s scoring system (Figure 4A). Our results showed that ZLN005 significantly elevates the latency to statge-4 seizures and reduces the average seizure scores in *Fmr1* KO mice with no apparent effects in WT mice. We also observed similar effects in vitro in cultured WT and *Fmr1* KO cortical neurons on a multielectrode array (MEA) recording system treated with ZLN005 (5 μM) or DMSO for 24 hours. As shown in Figure 4B, ZLN005 significantly reduces total spike numbers and spike rate in *Fmr1* KO cultures with no effects in WT cultures. Burst effects (burst frequency, burst duration and numbers of spike per burst) and synchrony index were not affected by ZLN005 in either WT or *Fmr1* KO cultures. Together, these data suggest that ZLN005 reduces hyperexcitability phenotypes in *Fmr1* KO mice.

**Figure 4.**
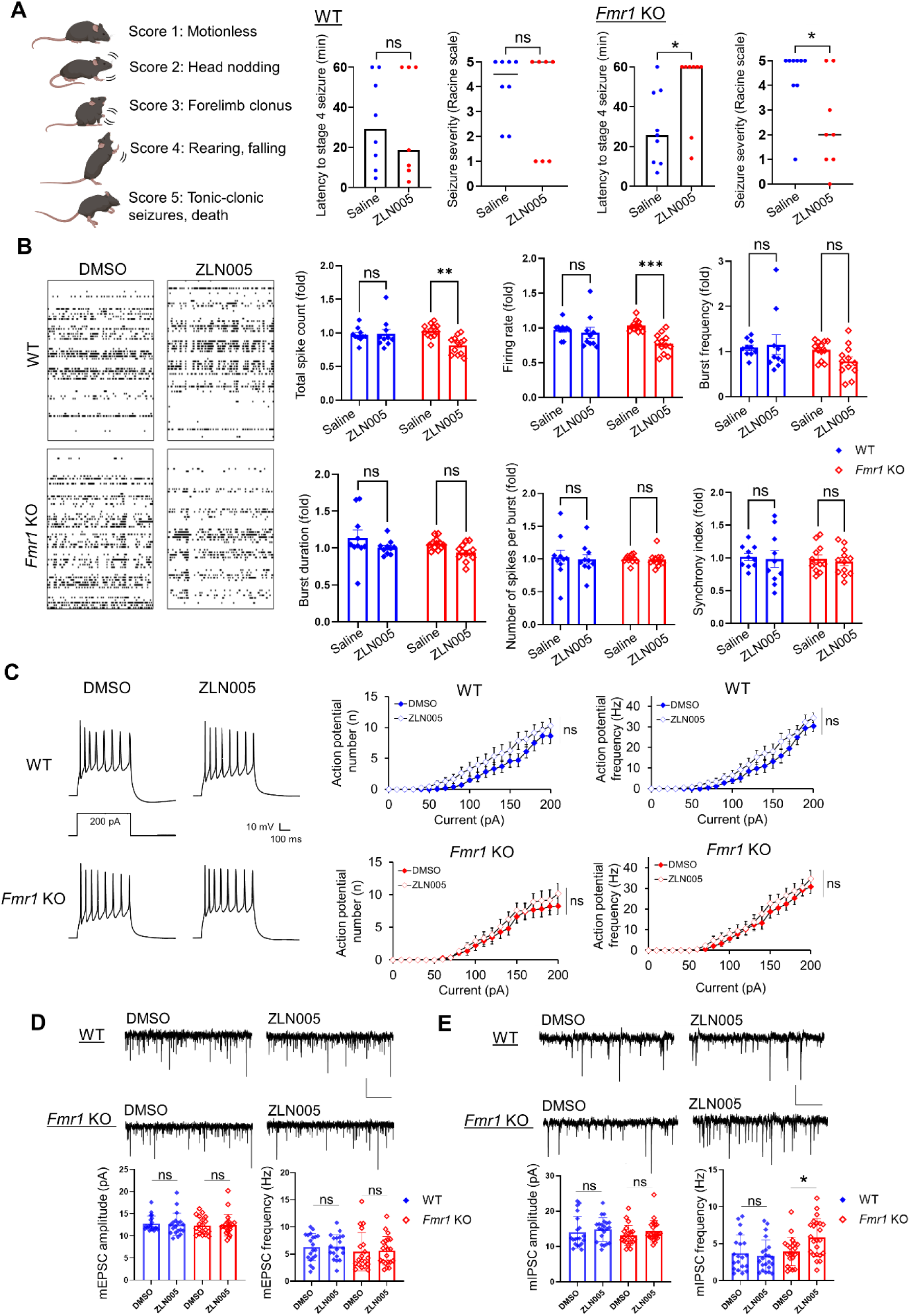
ZLN005 ameliorates hyperexcitability phenotypes in *Fmr1* KO mice. (**A**) Left: Racine scale schematic for seizure scoring. Right: Effect of ZLN005 on kainic acid (30 mg/kg, ip)–induced status epilepticus in WT (n = 7–8) and *Fmr1* KO (n = 8–9) mice following daily saline or ZLN005 treatment (2.5 mg/kg, i.p.) for 7 days. Latency to stage 4 was set to 60 min if seizures did not reach stage 4. (**B**) Representative MEA raster plots of spontaneous spikes from WT (n = 3, top) and *Fmr1* KO (n = 4, bottom) cortical neuron cultures treated with DMSO or ZLN005 (5 µM) for 24 h at DIV 12–14. Right: Bar graphs showing ratios (post-/pre-treatment) of total spike count, firing rate, burst frequency, burst duration, number of spikes per burst, and synchrony index. (**C**) Left: Action potential traces induced by 200 pA in WT (top) and *Fmr1* KO (bottom) neurons treated with DMSO or ZLN005 (5 µM) for 24 h. Right: Average firing rates (Hz) evoked by 0–200 pA injections. (n = 14-15 neurons per group). (**D**) Representative traces of mEPSCs (top) and quantification (bottom) from WT and *Fmr1* KO neurons treated with DMSO or ZLN005 (5 µM) for 24 h. (n = 19-21 neurons per group). (**E**) Representative traces of mIPSCs (top) and quantification (bottom) from WT and *Fmr1* KO neurons treated with DMSO or ZLN005 (5 µM) for 24 h. (n = 20-23 neurons per group). Mann-Whitney test (A) and two-way ANOVA with posthoc Tukey test (B-E) was used to calculate statistical significance. Data are presented as mean± SEM. *p < 0.05, **p < 0.01, ***p < 0.005, ns: non-significant.

The findings above prompted us to evaluate the effect of ZLN005 at the cellular level. To this end, we performed whole-cell patch-clamp recording in cultured WT and *Fmr1* KO cortical neurons. We first used a current-clamp recording to measure the action potential firing rate after delivering constant somatic current pulses for durations of 500 ms in the range of 0 to 200 pA (*37*). As shown in Figure 4C, ZLN005 does not elicit any significant effects on either WT or *Fmr1* KO neurons, suggesting the intrinsic excitability is likely unaffected. We followed by measuring the miniature excitatory postsynaptic currents (mEPSCs) and inhibitory postsynaptic currents (mIPSCs) in cultured primary cortical neurons. As shown (Figure 4D), ZLN005 does not exert any effects on mEPSCs in either WT or *Fmr1* KO neurons. Importantly, ZLN005 significantly elevates mIPSCs in *Fmr1* KO neurons without apparent effects in WT neurons (Figure 4E). These data suggest that ZLN005 reduces hyperexcitability phenotypes in *Fmr1* KO potentially by promoting inhibitory synaptic transmission.

### ZLN005 improves hippocampal short-term plasticity in *Fmr1* KO mice

Altered hippocampal plasticity in FXS is associated with various behavioral deficits. To investigate how ZLN005 treatment affects hippocampal plasticity in vivo, we chose to measure short-term plasticity because previous work has observed exaggerated synaptic facilitation in hippocampal slices of *Fmr1* KO (*38*). To this end, we measured the slope of field excitatory post-synaptic potential (fEPSP) in anesthetized mice in vivo (Figure 5A). We first compared the basal synaptic transmission between WT and *Fmr1* KO mice and observed no significant difference in saline- (Figure S2) or ZLN005-treated mice (Figure 5B). The effect was not different in mice anesthetized by isoflurane or urethane (Figure S2). However, when we quantified paired-pulse ratios, we observed a significant increase in paired-pulse ratios in saline treated *Fmr1* KO mice compared to WT mice across all stimulation intensities (Figure S2), confirming an altered short-term plasticity in *Fmr1* KO mice as described before (*38*). When evaluating the effect following daily ZLN005 (2.5 mg/kg) treatment for 7 days, we observed significantly decreased paired-pulse ratio across all stimulation intensities in *Fmr1* KO mice with no effect in WT mice (Figure 5C). We further examined the effect of ZLN005 treatment on short-term plasticity by delivering paired pulses at different inter-stimulation intervals. Similar to above, ZLN005 treatment significantly decreased paired-pulse ratios in *Fmr1* KO mice and an interaction between treatment and inter-stimulation intervals was found (Figure 5D). As similarly observed in previous data sets, no effect of ZLN005 was found in WT mice. Together, our data suggest that ZLN005 rectifies hippocampal short-term plasticity in *Fmr1* KO mice with no apparent effects in WT mice.

**Figure 5.**
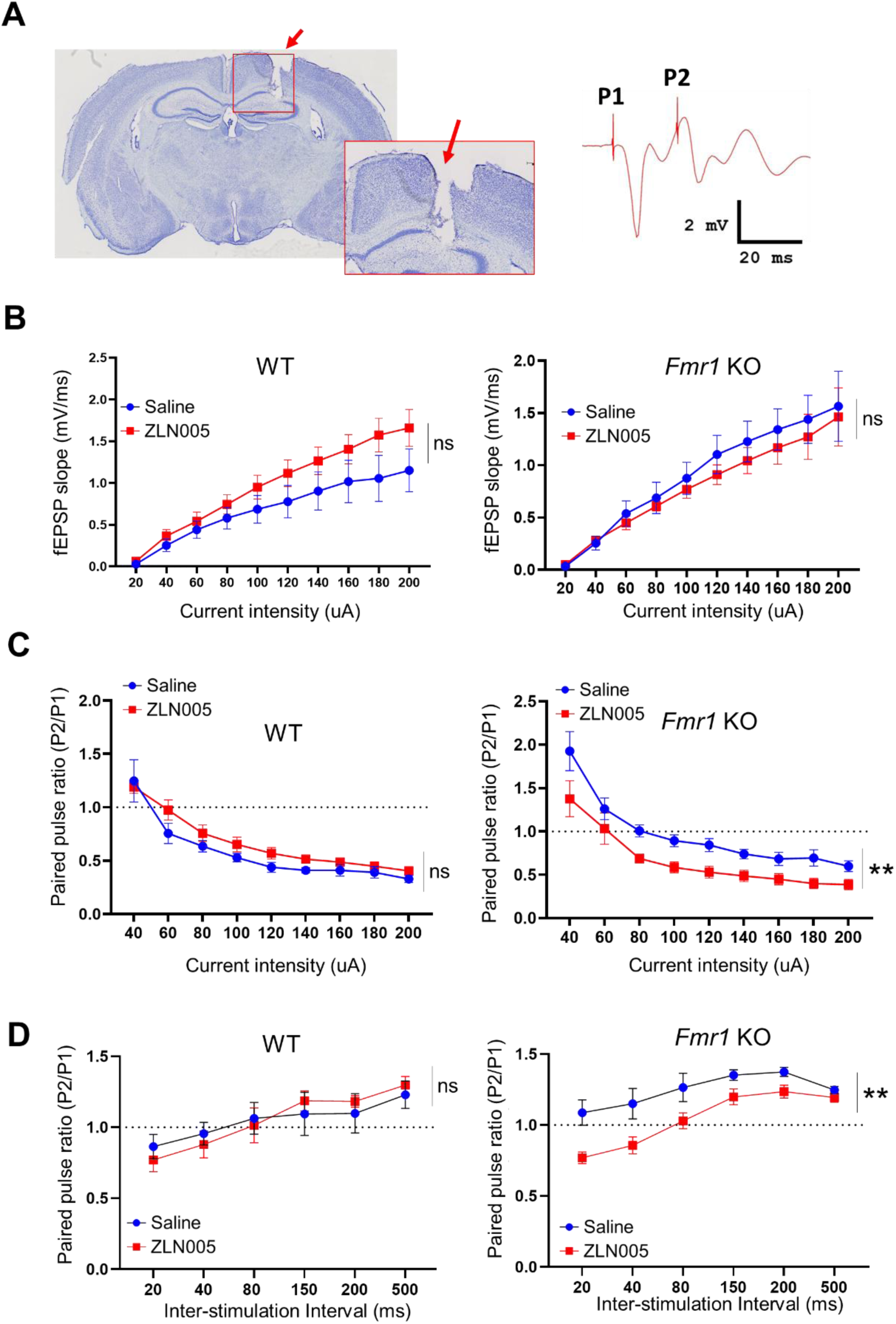
ZLN005 rectifies hippocampal short-term plasticity in *Fmr1* KO mice. (**A**) Representative cresyl violet–stained brain section showing the placement of the recording electrode in the hippocampal CA1 subfield. Field excitatory postsynaptic potentials (fEPSPs) were recorded by delivering paired electrical pulses (P1 and P2) in isoflurane-anesthetized mice. (**B**) Input–output curves of fEPSP slope for WT (n = 5) and *Fmr1* KO (n = 6-7) mice and the (**C**) paired-pulse ratios (PPR; P2/P1) measured 20-ms inter-stimulus interval following daily saline or ZLN005 treatment (2.5 mg/kg, i.p.) for 7 days in WT (n = 5) and *Fmr1* KO (n = 6-7) mice. (**D**) PPR of fEPSP slope measured across multiple inter-stimulus intervals in WT (n = 5) and *Fmr1* KO (n = 6-7) mice after daily saline or ZLN005 treatment (2.5 mg/kg, i.p.) for 7 days. Two-way ANOVA was used to calculate statistical significance. Data are presented as mean ± SEM. ** *p* < 0.01, ns: non-significant.

### ZLN005 ameliorates aberrant high-gamma band power and improves interhemispheric theta coherence in *Fmr1* KO mice

In EEG and LFP recordings, an increase in resting-state gamma-band power was reported in both human patients and animal models of FXS (*39–41*). Such an effect is thought to contribute to hyperexcitability in neuronal circuits and altered information processing in FXS (*42*). Our results corroborate these findings, as *Fmr1* KO mice showed an increase in both low-gamma (LG) and high-gamma (HG) band power in primary sensory cortex (S1) as well as in primary auditory cortex (Au1) when compared to wild-type (WT) mice (Figure 6A-6B). When we treated the mice with daily ZLN005 administration (2.5 mg/kg) for 7 days, despite no effect was found on LG band power in *Fmr1* KO mice, treatment with ZLN005 significantly reduced the aberrant increase in high-gamma band power in *Fmr1* KO mice in S1 (Figure 6A) and Au1 (Figure 6B). As we observed in previous data sets, no effect of ZLN005 treatment was found in WT mice. Additionally, we also found that ZLN005 treatment improved interhemispheric theta-band and alpha-band coherence (S1 – Au1) in *Fmr1* KO mice (Figure 6C). No significant effect of ZLN005 treatment was found in WT mice. Altogether, our data showed that ZLN005 ameliorates HG band activity and improved interhemispheric coherence in *Fmr1* KO mice.

**Figure 6.**
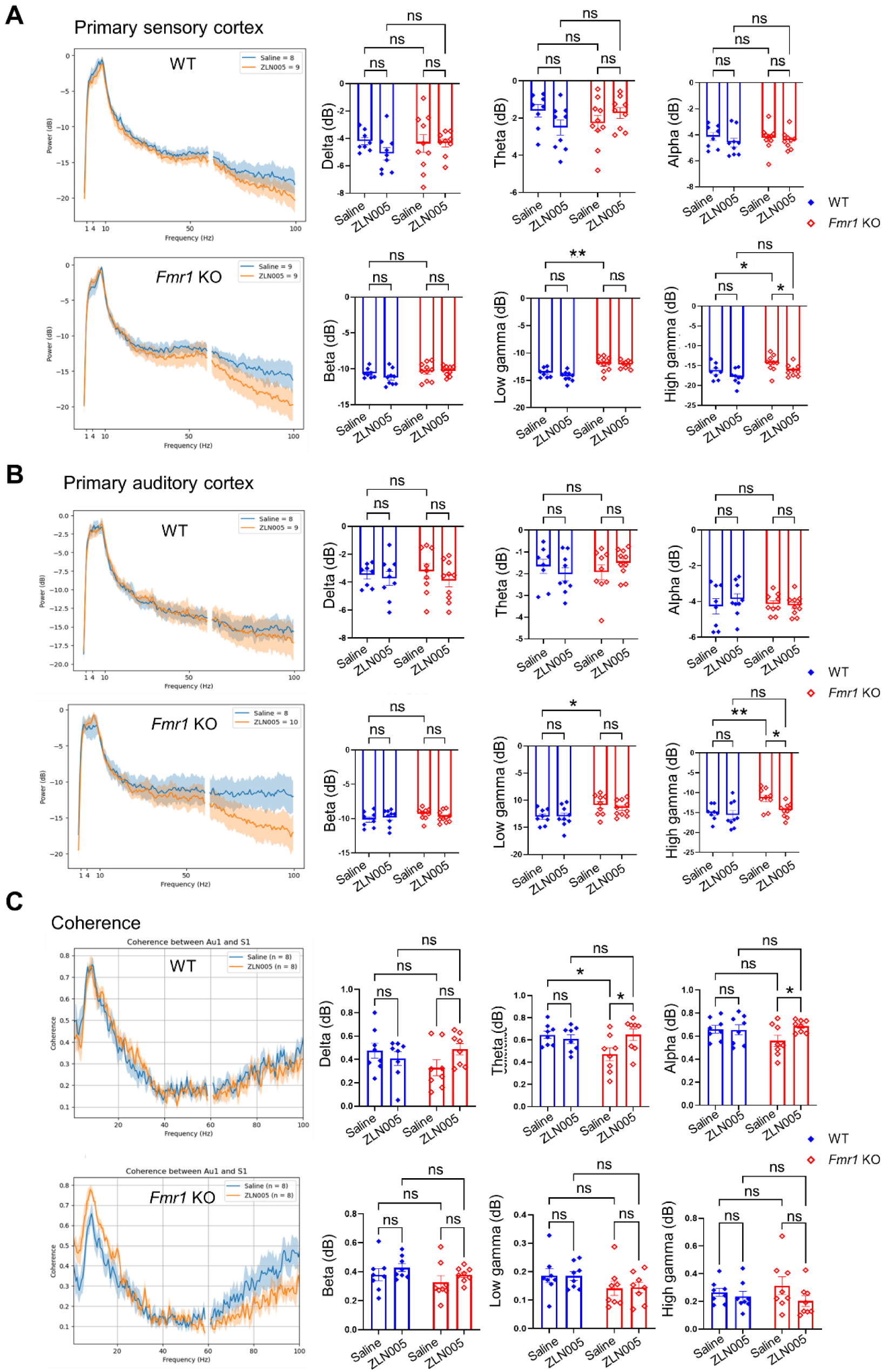
ZLN005 ameliorates high-gamma band power in *Fmr1* KO mice. (**A, B**) Power spectral density plots from (A) primary sensory and (B) primary auditory cortices of WT (n = 8-9) and *Fmr1* KO (n = 9-10) mice following daily saline or ZLN005 treatment (2.5 mg/kg, i.p.) for 7 days. The data in the spectral plots represents mean ± 95% CI half-width. The data in bar graphs for various frequency bands are represented as mean ± SEM. (**C**) Baseline coherence between primary sensory and primary auditory cortices from WT (n = 8-9) and *Fmr1* KO (n = 9-10) mice following daily saline or ZLN005 treatment (2.5 mg/kg, i.p.) for 7 days. Two-way ANOVA was used to calculate statistical significance. The data is represented as mean ± SEM in both line graphs and bar graphs. * *p* < 0.05, ** *p* < 0.01, ns: non-significant.

### ZLN005 improves cognition and reduces autism-like phenotypes in *Fmr1* KO mice

We next asked how ZLN005 may impact on cognition and autism-like behavior in *Fmr1* KO mice. We first performed an open field test to examine locomotor activity and anxiety-like behavior. As shown in Figure 7A, both WT and *Fmr1* KO mice receiving either daily saline or daily ZLN005 (2.5 mg/kg) treatment displayed similar levels of distance traveled, average speed, immobile time, and time spent in a center zone. These data suggest that ZLN005 does not affect locomotor activity and anxiety-like behavior in mice. We followed by employing a three-chamber social interaction test to measure the sociability behavior, which is often impaired in autism. After allowing the mice to habituate to the chamber as well as the empty wired cylinders, a stranger mouse was put into one of the cylinders and the time spent in the right and left compartments was measured. As shown in Figure 7B, both WT and *Fmr1* KO mice receiving either saline or ZLN005 spent significantly more time with the stranger mouse than with the empty cylinder. No drug or genotype difference was observed. These data suggest that *Fmr1* KO exhibit no significant defects in sociability and ZLN005 does not affect this behavior. To further probe autism-related behavior, we employed a marble burying test to evaluate repetitive behaviors, which are also an autism-linked behavior. As shown in Figure 7C, ZLN005 significantly reduces the number of marbles buried by *Fmr1* KO while no significant effects were observed in WT mice. These data suggest that ZLN005 reduces repetitive behavior in *Fmr1* KO mice.

**Figure 7.**
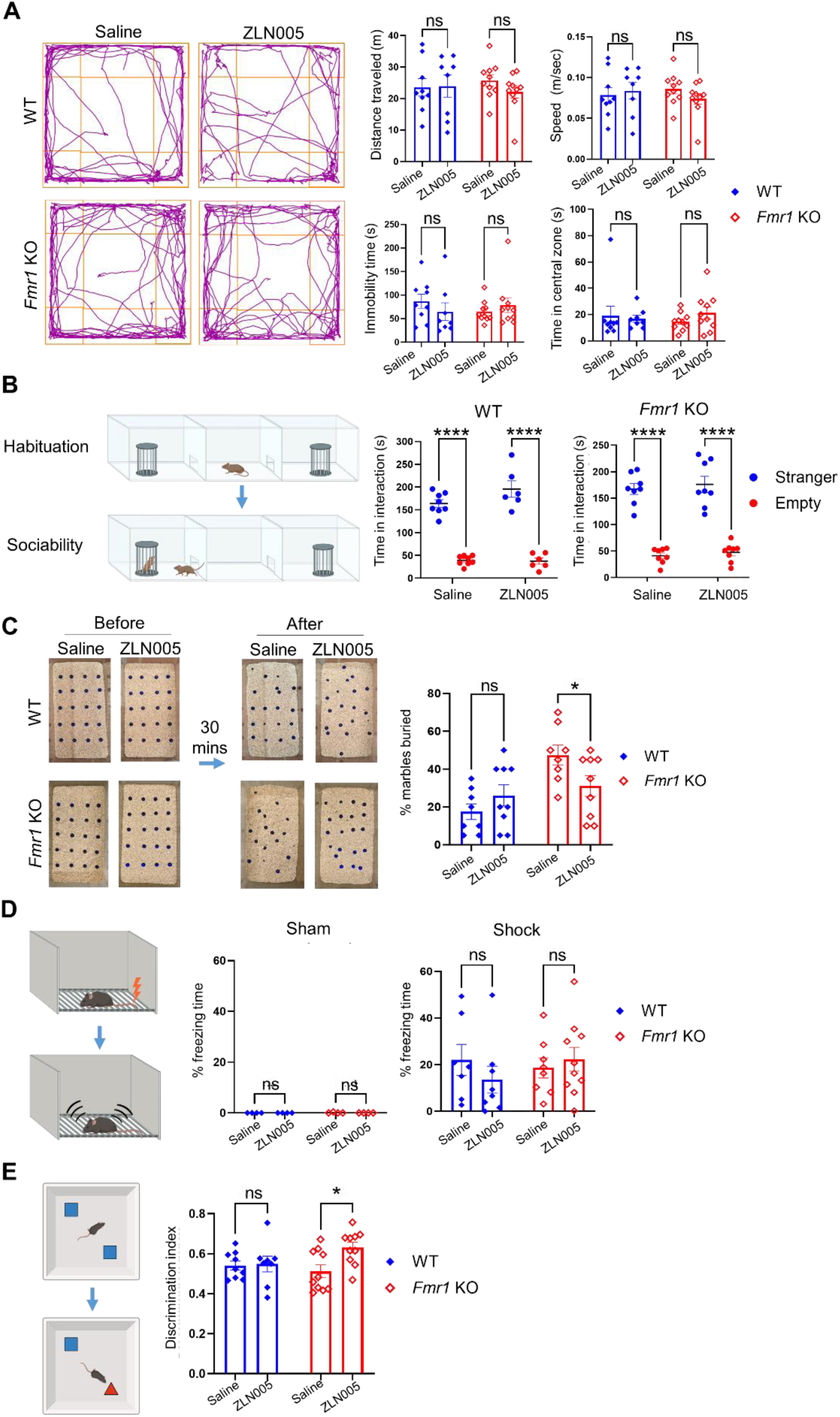
ZLN005 improves cognition and reduces autism-like behavior in *Fmr1* KO mice. (**A**) Representative traces from a 5-min open-field test (left) showing movement of WT (n = 8-9) and *Fmr1* KO (n = 10) mice after daily saline or ZLN005 treatment (2.5 mg/kg, i.p.) for 7 days. Quantification of distance traveled, speed, immobility time, and time spent in the center is shown on the right. (**B**) Three-chamber social interaction test for WT and *Fmr1* KO mice after daily saline or ZLN005 treatment (2.5 mg/kg, i.p.) for 7 days, with schematic of the sociability protocol on the left (n = 6-8). (**C**) Images of marbles before and after 30-min interaction (left) and quantification of marble-burying activity (right) in WT and *Fmr1* KO mice after daily saline or ZLN005 treatment (2.5 mg/kg, i.p.) for 7 days (n = 8-9). (**D**) Schematic of contextual fear conditioning (left) and freezing behavior quantification (right) in WT (n = 7-8) and *Fmr1* KO (8–10) mice after daily saline or ZLN005 treatment (2.5 mg/kg, i.p.) for 7 days. (**E**) Schematic of novel object recognition test (left) and preference index quantification (right) in WT (n = 8-9) and *Fmr1* KO (n = 10) mice after daily saline or ZLN005 treatment (2.5 mg/kg, i.p.) for 7 days. Two-way ANOVA was used to calculate statistical significance. Data are presented as mean ± SEM; * *p* < 0.05, ** *p* < 0.01, ns: non-significant.

To assess the effects of ZLN005 on cognition, we first employed the contextual fear conditioning test to test associative learning and memory. During the training day, mice received brief aversive stimuli (footshock, 2×, 0.5 mA, 2 s duration, 30 s interval). On the test day, mice were re-exposed to the same training apparatus and assessed for fear memory by measuring freezing behavior in the absence of footshock stimuli. As shown in Figure 7D, both WT and *Fmr1* KO mice receiving either saline or ZLN005 exhibit similar freezer behavior, indicating no significant effects on their associative learning and memory. We next employed a novel object recognition test to test object recognition memory. WT and *Fmr1* KO mice injected with saline or ZLN005 were allowed to explore two identical objects during the first session, followed by a second session where one of the two original objects was replaced with a novel object. As shown in Figure 7E, in comparison to WT mice, *Fmr1* KO mice exhibited an induction in recognition memory by ZLN005, demonstrated by a significant elevation in preference index. Altogether, our results suggest that ZLN005 reduces selective autism-like behavior and enhances cognition in *Fmr1* KO mice with no apparent effects in WT mice.

## DISCUSSION

In this study, we showed that a PGC-1α activator, ZLN005, promotes mitochondrial biogenesis and reverses many common pathophysiological defects in *Fmr1* KO mice. We also revealed that this effect is through AMPK-dependent phosphorylation of CREB. To our knowledge, this is the first evidence demonstrating the mechanism by which ZLN005 promotes PGC-1α expression. Most importantly, we showed that these effects are *Fmr1* KO-dependent likely because CREB phosphorylation is basally reduced in *Fmr1* KO mice, rendering them more responsive to ZLN005-induced CREB phosphorylation. A reduction of cyclic AMP (cAMP) signaling, which was seen in FXS (*43*), could be responsible for CREB dephosphorylation in *Fmr1* KO mice. Promoting cAMP production through inhibition of phosphodiesterases has shown promises in reducing FXS phenotypes (*44, 45*). Based on our current findings, it is possible that the beneficial effects of promoting cAMP production in FXS, as seen in prior studies, result in part from elevated CREB phosphorylation, leading to enhanced PGC-1α expression and mitochondria biogenesis. The potential crosstalk between cAMP signaling and mitochondria in FXS necessitates future experiments to explore and validate.

Many targets of FMRP encode mitochondrial proteins, and, as a result, loss of FMRP leads to profound mitochondrial defects observed in different animal and cell models. For example, impaired mitochondrial membrane integrity and activity were observed in the mouse model of FXS (*8, 46*). In the Drosophila model of FXS, altered mitochondria-endoplasmic reticulum contact sites and energy metabolism were detected (*9, 47*). In human FXS patient stem cell-derived neurons, changes in mitochondrial morphology and size were seen (*11*). Although the findings are variable, the consensus is that boosting mitochondrial activity appears to reverse FXS phenotypes. Our findings are in line with previous research and suggest PGC-1α as a major driving force to reverse mitochondrial defects and pathophysiology in the FXS mouse model. Our data also indicate that other chemicals that promote PGC-1α may be also beneficial to FXS. For example, a naturally occurring polyphenol named Resveratrol can promote mitochondrial functions and energy metabolism by promoting deacetylation of PGC-1α and thereby activating it (*48*). Resveratrol, or other drugs that promote AMPK, CREB phosphorylation, and PGC-1α activation, may boost mitochondrial functions and ameliorate FXS phenotypes. We believe this is a promising direction to search for or develop future FXS therapies.

In animal models and human patients with FXS, an increase in cortical EEG/LFP gamma band power (30-100 Hz) has been reported (*39–41*). The cortical low-gamma band (30 – 60 Hz) power reflects the interaction between pyramidal neurons and parvalbumin interneurons (*49*) while the high-gamma band (60 – 100 Hz) power is attributed to local neuronal spiking activity (*50, 51*).

The aberrant increase in high gamma band power during FXS is resultant of lower firing thresholds, increased firing frequency, and bursting activity of neurons (*52, 53*). This hyperexcitation may require higher energy production, consequently altering mitochondrial biogenesis and functions in FXS. Our data show that ZLN005 treatment improves mitochondrial functions and reduces high-gamma band power in freely behaving *Fmr1* KO mice during LFP recording. Because the increase in high-gamma band power in FXS could act as a background noise and reduce the signal to noise ratio (*51*), reducing high-gamma band power likely enables brain regions to coordinate better while improving sensory integration. This observation is further supported by our data showing significant improvement in selective behavioral tests in ZLN005-treated *Fmr1* KO mice.

Our whole-cell patch-clamp experiments show that ZLN005 significantly increased mIPSC frequency only in *Fmr1* KO neurons, suggesting a genotype-specific effect on inhibitory synaptic activity. These observations align with prior evidence that loss of FMRP impairs multiple aspects of GABAergic signaling, including phasic and tonic inhibition (*54*), GABA_A_ receptor-mediated signaling (*51*), delayed GABAergic neurogenesis (*55*), and an excitatory/inhibitory imbalance (*56*). Consequently, interventions that enhance inhibitory tone have consistently improved cognitive and behavioral outcomes in animal models of FXS, reinforcing the therapeutic relevance of targeting GABAergic pathways (*57–59*). The same concept can also explain our observation in hippocampal short-term plasticity. Our data revealed an increase in paired pulse ratio in *Fmr1* KO mice in vivo when compared to wildtype mice at a shorter inter-stimulation interval of 20 ms. A decrease in paired pulse ratio at shorter inter-stimulation intervals typically occurs due to GABA_A_-mediated feedback inhibition of CA1 pyramidal neurons (*60*). In *Fmr1* KO mice, treatment with ZLN005 restored paired pulse ratio to wildtype levels, further supporting the hypothesis that an increase in PGC-1α expression likely promotes GABAergic neurotransmission in *Fmr1* KO mice by either presynaptically increasing GABA release or postsynaptically by regulating GABA_A_ receptor expression (*61, 62*). Altogether, our findings suggest a role of ZLN005 in reshaping functional circuit activity and connectivity in *Fmr1* KO mice likely through facilitating inhibitory synaptic transmission.

## MATERIALS AND METHODS

### Study Design

To minimize the variability, sister cultures made from the same litter of at least three mice were prepared at the same time. The cell suspensions were pooled before plating. For western blotting, at least three different litters were used for each experiment. For seizure experiments, at least three litters of littermates were used for each condition in each experiment. For immunocytochemistry, at least two litters were used for each experiment. No randomization was performed to allocate subjects in this study. No exclusion criteria were predetermined and no mice were excluded. Because of the design of our research, no blinding was performed. Expected effect sizes and sample sizes for experiments were decided based on our previous studies (*63–66*).

### Statistical Analysis

The data presented in this study have been tested for normality using the Kolmogorov-Smirnov test. Student’s *t*-test was used when two conditions or groups were compared. Two-tailed Mann-Whitney test was performed as non-parametric test to compare two groups when the criteria for Student’s t-test were not met. Two-way ANOVA or mixed-effects model with Geisser-Greenhouse correction was used to compare the data with genotype and treatment as factors. For *post hoc,* Holm-Šidák*’*s test or Fisher’s LSD test was used when making multiple comparisons.

Due to the design of our research, no blinding was performed. Specific sample numbers, including the numbers of cells or repeats, are indicated in the figure legends. Data analyses were performed using GraphPad Prism software. Differences are considered significant when *p* < 0.05.

### Animals

All experiments using animals followed the guidelines of Animal Care and Use provided by the Illinois Institutional Animal Care and Use Committee (IACUC) and the guidelines of the Euthanasia of Animals provided by the American Veterinary Medical Association (AVMA) to minimize animal suffering and the number of animals used. This study was performed under an approved IACUC animal protocol of University of Illinois at Urbana-Champaign (23016 to N.-P. Tsai.). We obtained WT (stock No. 00664) and *Fmr1* KO mice (stock No. 003025) from Jackson laboratory. Because of the higher prevalence of FXS in males, we only employed male mice in our study. Mice were housed in individually ventilated cages in 12 hrs light/dark cycle with *ad libitum* access to pelleted food and water.

### Serial block**-**face scanning electron microscopy (SBF-SEM)

Mice treated with saline or ZLN005 were anesthetized with isoflurane and perfused transcardially with 4% glutaraldehyde and 2% paraformaldehyde in 0.1 M cacodylate buffer (pH 7.4, 2 mM CaCl_2_Ted Pella Inc.) Brains were then isolated and stored in fixative at 4°C overnight. Fixed brains were washed 5 times, 3 minutes each, with ice-cold cacodylate buffer (0.15 M cacodylate buffer with 2 mM CaCl_2_) to remove the fixative, sectioned on a vibratome in ice-cold cacodylate buffer, and 200-µm sections were obtained. The brain sections were then sequentially incubated in 2% osmium tetroxide (OsO_4_; Electron Microscopy Sciences) prepared in 0.15 M cacodylate buffer for 90 min at room temperature (RT), then transferred without washing to 2.5% potassium ferrocyanide (Sigma-Aldrich) prepared in the same buffer (90 min at RT) to form a reduced osmium complex. The sections were then washed twice in ddH_2_O for 30 min each and incubated in 1% thiocarbohydrazide (TCH; Sigma-Aldrich) for 45 min at 40°C. Sections were washed twice as above and re-incubated in 2% OsO_4_ (90 min at RT), followed by two final washes with ddH_2_O. En bloc staining was then performed by incubating sections in 1% uranyl acetate (Electron Microscopy Science) (overnight at 4°C and then 2 h at 50°C) and freshly prepared lead aspartate solution (pH 5.0; 2 h at 50 °C) with twice ddH_2_O washes after each step. After the staining, the sections were dehydrated through an ethanol gradient (50%, 75%, 100%; 30 min each, on ice) and then transferred to ice-cold anhydrous acetone (3 changes, 30 min each at RT). Sections were infiltrated stepwise with Durcupan ACM resin in acetone (25%, 50%, 75%, and twice at 100%, 2 h each; 100% resin overnight), and sections were finally polymerized at 60°C for 48 hr. For imaging, the cured blocks were trimmed (≤1 mm³–2 mm × 2 mm × 1 mm), mounted on Gatan 3View aluminum pins with conductive silver epoxy (Circuit Works, CW2400), and edge-coated with silver paint or an AuPd sputter coat. The SBF-SEM imaging was performed on a Zeiss Sigma VP field-emission SEM with a Gatan 3View2XP ultramicrotome at 2 kV, dwell time 1 μs, with focal charge compensation, acquiring 16,000 × 16,000-pixel images at 1-nm x–y pixel size and 50 nm z-axis slice thickness.

### SBF-SEM image segmentation and deep learning–based analysis

We created in-house models to segment mitochondria and nuclei present within our SBF-SEM data. For both models, we utilized a machine-learning U-network with attention features, with the full model architecture (Figure S1A) (https://github.com/vipendrakumarsingh/SBF-SEM-image-segmentation). Previous work has thoroughly shown that this type of model is uniquely suited for tasks such as image segmentation (*67*). The model works by downsizing the original input image through convolutional and pooling layers until it reaches the center point, termed the bottleneck. The model then runs through further convolutional and up-sampling layers till it returns an image that is the same size as the input. The image that is generated is a binary mask of the organelle of interest. These binary masks contain a 0 or 1 depending on whether the model determines that the organelle of interest is absent or present at the specified pixel. The model architecture is uniquely suited for this task, since the bottleneck forces information fed through to retain only the most vital components that pertain to generating the binary mask. Additionally, our model contains convolutional block attention modules that serve to refine feature extraction within each layer.

To train the model, mitochondria or nuclei within SBF-SEM image slices were first annotated manually using the Webknossos web service. These annotations were saved and reformatted into binary images. The binary images were then reformatted into multiple non-overlapping 200 × 200 pixel images. Using this method, we obtained 1175 images, of which 1000 served as input to train the model and 175 as test images (batch size = 4, epochs = 1000; training curves are provided in Figure S1C). Once the model is trained, we annotate full SBF-SEM image slices via an inference function that takes patches of the image to pass through the model. Patches are subsequently combined to generate a full image mask with the same dimensions as the original image. Masks for the cell bodies were performed manually via the microscope image browser application.

### 3D reconstruction and analysis

Binary masks corresponding to mitochondria and nucleus generated by our attention-UNET model, and the cell body model generated using the Microscopy Image Browser (MIB). These masks were imported into AMIRA for 3D reconstruction and analysis. The mitochondria, nucleus, and cell body in each layer were tracked, and label analysis was performed to create label fields for quantification of morphometric data, such as volume and surface area, and generate a 3D model for visualization. To limit mitochondrial measurements to the cytoplasm of the segmented cell body, a logical masking operation was applied in Amira using the expression Ax(B-C), where A is the mitochondrial mask, B is the cell body mask, and C is the nuclear mask. Next, the mitochondrial volume and surface area of all mitochondria were exported to a CSV file, and thresholding was applied to minimize segmentation noise, boundary artifacts, and partial fragments. Objects smaller than 0.0003 µm^3^ (approximately 250 voxels) and surface area smaller than 0.02 µm^2^ were excluded. Morphometric data were then exported and used to calculate log-transformed volume, mitochondrial complexity index (MCI), and sphericity. The following formula was used to calculate the MCI as utilized before (*68*): MCI=SA^3^/16π^2^V^2^ where SA is surface area, and V is volume of mitochondria. The sphericity of the mitochondria was calculated using the formula Ψ=(π^1/3^(6V)^2/3^)/SA, where Ψ is the sphericity index, SA is the surface area, and V is the volume of mitochondria.

## List of Supplementary Materials

Supplemental Materials and Methods

Figures S1 and S2

Movie S1

## ACKNOWLEDGEMENTS

The authors would like to thank Dr. Catherine Christian-Hinman (University of Illinois at Urbana-Champaign) for providing technical support when setting up the EEG system.

## AUTHOR CONTRIBUTIONS

Conceptualization: AA, VK, N-PT Methodology: AA, VK, GG, KAB, AJC, JSR Investigation: AA, VK, LKY, MSB

Funding acquisition: AA, VK, N-PT Project administration: N-PT Supervision: N-PT

Writing: AA, VK, N-PT

## DATA AND MATERIALS AVAILABILITY

All data are available in the main text or the supplementary materials.

## SUPPLEMENTAL MATERIALS AND METHODS

### Primary neuronal culture

The primary cortical neuronal culture was performed using mice at post-natal day 0-1. Cortices were dissected and incubated with trypsin for 8-10 mins at 37°C. Next, trypsin was neutralized by the addition of fetal bovine serum (FBS) supplemented HBSS and washed twice with pre-warm HBSS. Cortices were then homogenized in complete DMEM and plated on poly-D-lysine (0.05 mg/ml) (sc-136156, Santa Cruz Biotechnology) coated 6-well plates. After 3-5 hrs, the medium was replaced with Neurobasal A medium (10888022, ThermoFisher Scientific) supplemented with 2 mM Glutamax (35050061, Invitrogen), B27 supplement (17504001, Invitrogen) and 1 µM (β-D-Arabinofuranosyl cytosine; Ara-C) (C1768, Sigma-Aldrich). Cultures were maintained at 37°C with 5% CO2. Half of the medium was changed on days-in-vitro (DIV) 2 and thereafter every 3-4 days. Experiments were performed when cultures were at DIV 14-16.

### Reagents

Dimethyl sulfoxide (#BP231) was from Thermo Fisher Scientific. ZLN005 (#14121) and BAY-3827 (#41448) were from Cayman Chemical. 666-15 (#GC32689) was from GlpBio. BAY-Antibodies for western blotting were purchased from Proteintech: mouse anti-GAPDH (#60004-1) and mouse anti-MFN2 (#67487-1); from Cell Signaling Technology: mouse anti-β-actin (#3700), rabbit anti-CREB (#9197), rabbit anti-phospho-CREB (#9198), rabbit anti-AMPK (#5831), rabbit anti-phospho-AMPK (#2535), rabbit anti-VDAC (#4661), rabbit anti-TOM20 (#4240), rabbit anti-NRF1 (#46743), mouse anti-COX IV (#11967) and rabbit anti-DRP1 (#8570); and from Santa Cruz Biotechnology: anti-PGC-1α (#sc-517380). Secondary antibodies were from Cell Signaling Technology: anti-mouse HRP (#7076) and from Jackson ImmunoResearch: anti-rabbit HRP (#711-035-152).

### Western blotting

Tissue or cell cultures were lysed in ice-cold lysis buffer (137mM NaCl, 20 mM Tris-HCL, 2mM EDTA and 1% Triton X-100, pH 8.0) supplemented with protease inhibitors (A32963, ThermoFisher Scientific) and phosphatase inhibitors (P2850; Sigma-Aldrich). Lysates were sonicated and centrifuged. Supernatants were collected and protein concentration was measured using Bradford’s method. SDS buffer (40% glycerol; 240 mM Tris-HCl, pH 6.8; 8% sodium dodecyl sulfate; 0.04% bromophenol blue; and 5% β-mercaptoethanol) was added to the lysates and heated at 95°C in a heating block for 10 mins. Samples were then separated on SDS-PAGE gel and transferred onto a PVDF membrane (sc-3723, Santa Cruz Biotechnology). Membranes were blocked with 1% bovine serum albumin (BSA, BP9706100, ThermoFisher Scientific) in Tris-buffered saline Tween-20 buffer (TBST; [20 mM Tris, pH 7.5; 150 mM NaCl; 0.1% Tween-20]) for 30 mins. Subsequently, membranes were incubated with primary antibodies overnight at 4°C. Next, the membranes were washed 3 times in TBST and incubated with HRP-conjugated secondary antibody in 5% non-fat skimmed milk in TBST for an hour at 25°C. Membranes were washed with TBST 3 times and developed by using an enhanced chemiluminescence reagent and detected by an iBright imaging system (ThermoFisher Scientific, Waltham, MA). Band of the protein of interest were analyzed by ImageJ software (National Institute of Health).

### Hippocampal evoked potential recordings in anesthetized mice

The mice were anesthetized with isoflurane (5% for induction and 1.5% for maintenance, 0.5 L/min medical O₂). Body temperature was maintained at 36 ± 1°C using a heating pad with a rectal probe for temperature feedback. Preoperatively, carprofen (5 mg/kg) was administered subcutaneously to provide analgesia. The mice were then placed in a stereotactic frame (RWD Life Science Co., LTD, China), the fur on the head was shaved, and a single incision was made to expose the skull. After clearing the periosteum with 3% hydrogen peroxide, the bregma was identified, and a burr hole of approximately 1.4 mm in diameter (AP: −2.0 mm, ML: +1.7 mm) was made using an electric micro-drill to implant bipolar stimulation and monopolar recording electrodes (polyimide-coated stainless-steel wire, 70 µm diameter; California Fine Wire, Grover Beach, CA, USA) in the hippocampus. Another burr hole was made anterior to bregma in the left frontal bone using a Dremel drill and a threaded drill bit of 0.9 mm diameter to anchor the ground electrode. The electrodes were then slowly lowered into the brain at the following coordinates: Recording—AP: −1.9 mm, ML: +1.4 mm, DV: −1.2 to −1.4 mm; Stimulation—AP: −2.0 mm, ML: +2.0 mm, DV: −1.4 to −1.6 mm, targeting the stratum radiatum of the hippocampus (Franklin and Paxinos, 2007). Biphasic, charge-balanced electrical pulses (200 µs) were used to elicit evoked potentials (EPs). The depth of the recording electrodes was adjusted based on auditory feedback of CA1 neuronal firing and further optimized until the electrode could acquire a field excitatory postsynaptic potential (fEPSP) with a negative potential. After 30 minutes, input–output (IO) relationships (basal synaptic transmission) were determined by applying paired pulses (interstimulus interval [ISI] of 20 ms) with increasing current intensities (20–200 μA in steps of 20 μA) every 30 s. This procedure was repeated four times. Next, the effect of treatment on short-term plasticity was examined. For each mouse, the stimulation intensity that produced approximately 40% of the fEPSP slope was selected from the IO recording. Using this stimulation intensity, paired pulses with varying ISIs (20, 40, 80, 150, 200, 500 ms) were delivered every 30 s. The ratio of the fEPSP slope of the second pulse to the first pulse (P2/P1) was calculated.

After the recordings, the mice were transcardially perfused with PBS followed by 4% PFA, and electrode placement was verified using cresyl violet staining. The evoked potentials (EPs) were acquired using Digitimer’s NeuroLog System (UK). Signals were recorded with an NL100 AK AC headstage unity-gain preamplifier and amplified 500-fold using an NL104A AC Preamplifier. They were then subjected to an analog bandpass filter (0.1 Hz–50 kHz) and a notch filter (60 Hz) using an NL125/6 Band Pass Filter. Data acquisition and stimulation were performed using a CED Micro 1401 (input range: 5 V) and controlled with Signal software (Cambridge Electronic Design Ltd., England). CA1 EPs were sampled at 10 kHz with 16-bit resolution. Charge-balanced biphasic square-wave pulses (200 μs per phase) were used to obtain the EPs via a DS4 constant-current linear stimulus isolator (Digitimer, UK).

### In vivo local field potential recordings in awake and freely moving mice

The surgeries on mice were performed as described earlier. After locating the bregma, burr holes were made in five locations: the primary auditory cortex (Au1- AP: -2.5 mm, ML: -4.5 mm), primary sensory cortex (S1 - AP: -1.0 mm, ML: +3.0 mm), hippocampal CA1 subfield (CA1 - AP: -1.9 mm, ML: +1.4 mm), and cerebellum (AP: 6.25 mm, ML: ±2.0 mm). Epidural electrodes were custom-made, the stainless steel micromachine screws (#NAS721CE80-080, Swissturn USA Inc., USA) were soldered to PFA-coated silver wires of 254 µm diameter (A-M Systems, USA). These electrodes were gently fastened to all burr holes except at the CA1. A PFA-coated silver wire of 127 µm diameter was used as a depth electrode to record from the CA1 subfield of the hippocampus. The depth electrode was lowered 1.2 mm from the surface of the brain and fixed in position with dental UV cement. The epidural electrodes located on the cerebellum served as ground and reference. The differential contact of the CA1 electrode was pseudo-referenced to the epidural electrode located on the cerebellum. After electrode placement, headmounts were prepared by first layering the skull with dental cement (C&B Metabond) and then casting an additional layer of generic dental cement.

The mice were allowed to recover for one week, and treatments were administered for an additional week before the mice were connected for EEG recordings. They were allowed to habituate in the enclosure for one day prior to the recordings. Recordings were simultaneously performed on 3–5 mice between 13:00 and 18:00 h for 30 minutes in the presence of the experimenter to observe behavioral state. Periods when the mice were awake and still for at least one minute were noted and used for analysis.

We utilized 3-channel EEG systems (#8200-K1-SE3, Pinnacle Technology Inc., USA) to record the signals. Local field potentials (LFPs) were filtered using an analog high-pass filter at 1 Hz, and a digital cutoff was set at 100 Hz during recording. The LFPs were amplified 100-fold by the headstage amplifier and sampled at 1 kHz. The data conditioning and acquisition systems had a 16-bit resolution, and Sirenia acquisition software was used to record the LFPs. One-minute segments of local field potential (LFP) data were converted to tab-separated values (TSV) files and analyzed using Python. Power spectral density (PSD) and coherence were computed via the Welch method implemented in the SciPy library. A sliding window of 2048 samples and a Hanning taper were applied during the Fast Fourier Transform (FFT) computation. The PSD data were converted to the decibel scale (10 × log_10_) and normalized by dividing the PSD for each mouse by its maximum value, so that the peak power equaled 0 dB. The NumPy library’s plotting function was used to visualize PSD, representing the mean and 95% confidence intervals (95% CI) for each group.

Any bad channels or poor electrode contacts were omitted from the analyses by checking raw traces and computing PSD values for each mouse. Coherence was then calculated for primary sensory and primary auditory cortical electrodes in mice where both electrodes were active, using the SciPy library’s coherence function. The data were plotted as mean and standard error of the mean (SEM) for each group. For statistical analysis, the PSD data from Au1 and S1 were divided into six frequency ranges: Delta (1–4 Hz), Theta (4–8 Hz), Alpha (8–13 Hz), Beta (13–30 Hz), Low-gamma (30–59 Hz), and High-gamma (61–100 Hz). In case of CA1, theta range was computed from 4 – 12 Hz.

### Immunohistochemistry and imaging

Primary cortical neurons were made from WT mice at P0 or P1 as described previously on glass coverslips at a density of 1.5 × 10^5^ cells. At DIV 14, cells were incubated with MitoTracker Dye (Thermo Fisher, M7510) for 30 minutes at 37°C, followed by washing once with phosphate buffered saline (PBS), fixing with fixation buffer (4% paraformaldehyde and 4% sucrose in PBS) for 15 minutes, and then permeabilizing with 0.5% Triton-X-100 in PBS for 5 minutes at room temperature. Following fixation and permeabilization, the neurons were incubated overnight with primary antibodies in 1% BSA in PBS. Following the overnight incubation, the samples were washed three times for ten minutes each with PBS, incubated with Alexa Fluor conjugated secondary antibodies for two hours at room temperature in 1% BSA in PBS, and then were washed an additional three times for ten minutes each with PBS. The coverslips were then mounted onto glass slides. Images were obtained using a Zeiss LSM 700 confocal microscope with 40x magnification and three different laser lines (488, 555, and 633 nm). The pinhole was set to 1 airy unit.

### MEA recording

Multielectrode array (MEA) recordings were performed using Maestro Edge (Axion Biosystems) with Cytoview MEA6 plates (6-well plates). Field potentials were recorded at each electrode relative to the ground electrode with a sampling frequency of 12.5 kHz. Followed by 30 mins baseline recording (before), neurons were treated with indicated drugs for 24 hrs and recorded for another 30 minutes. To avoid the effect of change in physical movement on network activity, only the last 15 minutes of the recordings were used for data analysis. Axis Navigator version 3.3 software (Axion Biosystems) was used for spike extraction from raw electrical signals. After filtering the spike detector setting for each electrode was independently set at the threshold of ±6 standard deviation. Therefore, activity above the threshold was counted as a spike and included in data for analysis. The total number of spikes was normalized to the number of electrodes in each well. The average number of spikes was calculated and expressed as fold changes with respect to the control. For detection of burst, a minimum of 5 spikes with a maximum 100 ms spike interval was set for individual electrodes. Analysis of burst duration and burst frequency was performed using Axis Navigator version 3.3 software.

### Whole-cell patch clamp recordings

Whole-cell patch-clamp recordings were performed on dissociated cortical neurons cultured for 12–16 days in vitro (DIV12–16). Neurons were visually identified using an upright microscope (BX51WI, Olympus). Recording pipettes (4–6 MΩ) were pulled from borosilicate glass capillaries (1.5 mm outer diameter; Sutter Instruments, P-97) and filled with appropriate internal solutions depending on the recording configuration. All recordings were conducted at room temperature (23–25 °C). The external solution contained (in mM): 119 NaCl, 2.5 KCl, 4 CaCl₂, 4 MgCl₂, 1 NaH₂PO₄, 26 NaHCO₃, and 11 D-glucose, continuously bubbled with 95% O₂/5% CO₂ (pH 7.4, 310 mOsm). Cells were excluded if their resting membrane potential was above −45 mV. Series resistance was not compensated. Only cells with stable series resistance (<30 MΩ) were included in the analysis. Signals were amplified using a Multiclamp 700B amplifier, digitized at 10 kHz with a Digidata 1550B interface, and recorded using pClamp software (version 10.6; Molecular Devices). Data analysis was performed using Clampfit (version 10.6) and Mini Analysis Program (Synaptosoft).

To record action potentials (APs), whole-cell current-clamp recordings were performed using an internal solution containing (in mM): 130 K-gluconate, 6 KCl, 3 NaCl, 10 HEPES, 0.2 EGTA, 4 Mg-ATP, 0.4 Na-GTP, and 14 Tris-phosphocreatine (pH 7.25, 285 mOsm). To pharmacologically block fast synaptic transmission, 20 μM CNQX (Tocris, #1045), 200 μM DL-APV (Tocris, #3693), and 20 μM bicuculline (Tocris, #0131) were added to the external solution. Neurons were held at −60 mV in current-clamp mode, and a series of depolarizing current steps (0–200 pA, 500 ms duration) were injected to evoke action potentials. The firing frequency was determined as the reciprocal of the interspike interval between the first and second action potentials.

Miniature excitatory postsynaptic currents (mEPSCs) were recorded in voltage-clamp mode at −60 mV using the same external solution supplemented with 0.5 μM tetrodotoxin (TTX; Cayman Chemical, #14964) and 20 μM bicuculline to block voltage-gated sodium channels and GABA_A_ receptor-mediated currents, respectively. The internal solution contained (in mM): 130 K-gluconate, 6 KCl, 3 NaCl, 10 HEPES, 0.2 EGTA, 4 Mg-ATP, 0.4 Na-GTP, 14 Tris-phosphocreatine, and 2 QX-314 (pH 7.25, 285 mOsm).

Miniature inhibitory postsynaptic currents (mIPSCs) were recorded in voltage-clamp mode at −60 mV using a high-chloride internal solution containing (in mM): 68 K-gluconate, 68 KCl, 3 NaCl, 10 HEPES, 0.2 EGTA, 4 Mg-ATP, 0.4 Na-GTP, 14 Tris-phosphocreatine, and 2 QX-314 (pH 7.25, 285 mOsm). The external solution was supplemented with 0.5 μM TTX, 200 μM DL-APV, and 20 μM CNQX to block sodium channels, NMDA receptors, and AMPA receptors, respectively.

### Behavioral seizure assessment

Mice at 8 weeks of age were intraperitoneally injected with kainic acid (KA) prepared in saline (Cayman Chemical, USA) at 30 mg/kg to induce seizures. After injection, mice were monitored for an hour in real time. Modified Racine’s scoring system was used to assess the seizure intensity.

### Behavioral tests

All the behavioral tests were conducted on WT and *Fmr1* KO mice starting at 6 weeks of age. Mice were brought to the behavior testing room 30 minutes before the test and housed in their home cages. The room was dimly lit at 50 lux and low background noise (approx. 65 db). Behavioral apparatuses were thoroughly cleaned before and after every test session with 70% ethanol to avoid olfactory bias. Detailed procedures are provided below.

### Marble burying test

In a polycarbonate cage (26 × 48 × 20 cm) a total of 20 marbles were placed on the surface of bedding which was approximately 5 cm deep. Marbles were arranged in a 5×4 array. Mouse was introduced into the cage and allowed to remain inside for 30 minutes. Afterwards, the mouse was removed, a photograph of the cage floor was taken, and the total number of buried marbles was counted. A marble was classified as “buried” if two-thirds or more of it was concealed beneath the bedding. To carry out the next set of experiments, a clean cage filled with new bedding and marbles was used, following the same procedure as above.

### Open field test

The open field test was carried out to evaluate the locomotion and anxiety behavior of mice. The mouse was placed in the center of a plexiglass box (67 × 67 × 31 cm) and allowed to freely explore the arena for 5 minutes. The movement of the mouse was recorded with an overhead camera, and the video was analyzed using the AnimalTracker plugin in ImageJ (National Institute of Health). The open-field arena was virtually divided into central and outer zones. The movement trajectory, velocity, immobile time, and total distance travelled by the test animal were visualized and analyzed using ANY-maze software.

### Social interaction test

The social interaction test was conducted in a plexiglass chamber measuring 20 × 40 × 25 cm and was divided into 3 parts by transparent walls with small openings to allow free movement of test animals between all three compartments. The test consisted of 2 sessions: habituation and sociability. Each session lasted for 10 mins and was video recorded using an overhead camera. During the habituation session, the mouse was introduced to the middle chamber and allowed to explore all three chambers. For the sociability session, a stranger mouse of the same age and sex was placed in a wired cylinder and placed in the left chamber while an empty wired cylinder was placed in the right chamber. The test mouse was then reintroduced into the middle chamber and allowed to explore the left and the right chamber. The video from the second session was analyzed to calculate sociability. The time spent by the test mouse interacting with the stranger mouse over an empty cage was represented as sociability.

### Novel Object Recognition test

To test the object recognition memory, novel object recognition test was performed. On day-1 (trial day), the mouse was placed in the empty testing chamber (25 × 25 cm) and allowed to habituate for 10 min, during which the mouse was allowed to freely explore two identical objects in the box before returning to the home cage. On day 2 (testing day), one of the two identical objects was replaced with a novel object, and the mouse was allowed to explore them for 5 minutes. The preference for exploring the novel object was calculated by dividing the time elapsed by the mouse exploring the novel object (T_novel_) by the time elapsed for exploring both familiar and novel objects (T_novel_+T_familiar_) and expressed as the preference index i.e. (T_novel_/(T_novel_+T_familiar_) x100).

### Contextual fear conditioning

The test mice were divided into two groups: the shock (receiving foot shock) and the sham (not receiving foot shock). The experiment was carried out in two days. On the first day (training phase), the mice were placed inside the fear conditioning chamber (32 × 28 × 30 cm) with a metal grid floor for 3 minutes. The fear group received two foot shocks of 0.75 mA for 2 seconds at 120 sec and 150 sec. The control group was also placed in the box for 3 minutes but did not receive any foot shock. After 3 minutes, the test mice were returned to their home cage. On the second day, both the control group and the fear group were placed in the fear conditioning chamber for 3 mins. At this time no foot shock was delivered to either group. Each session was video-recorded and analyzed for freezing behavior. Freezing time was calculated and represented as the freezing percentage.

## SUPPLEMENTAL FIGURES AND FIGURE LEGENDS

**Figure S1.**
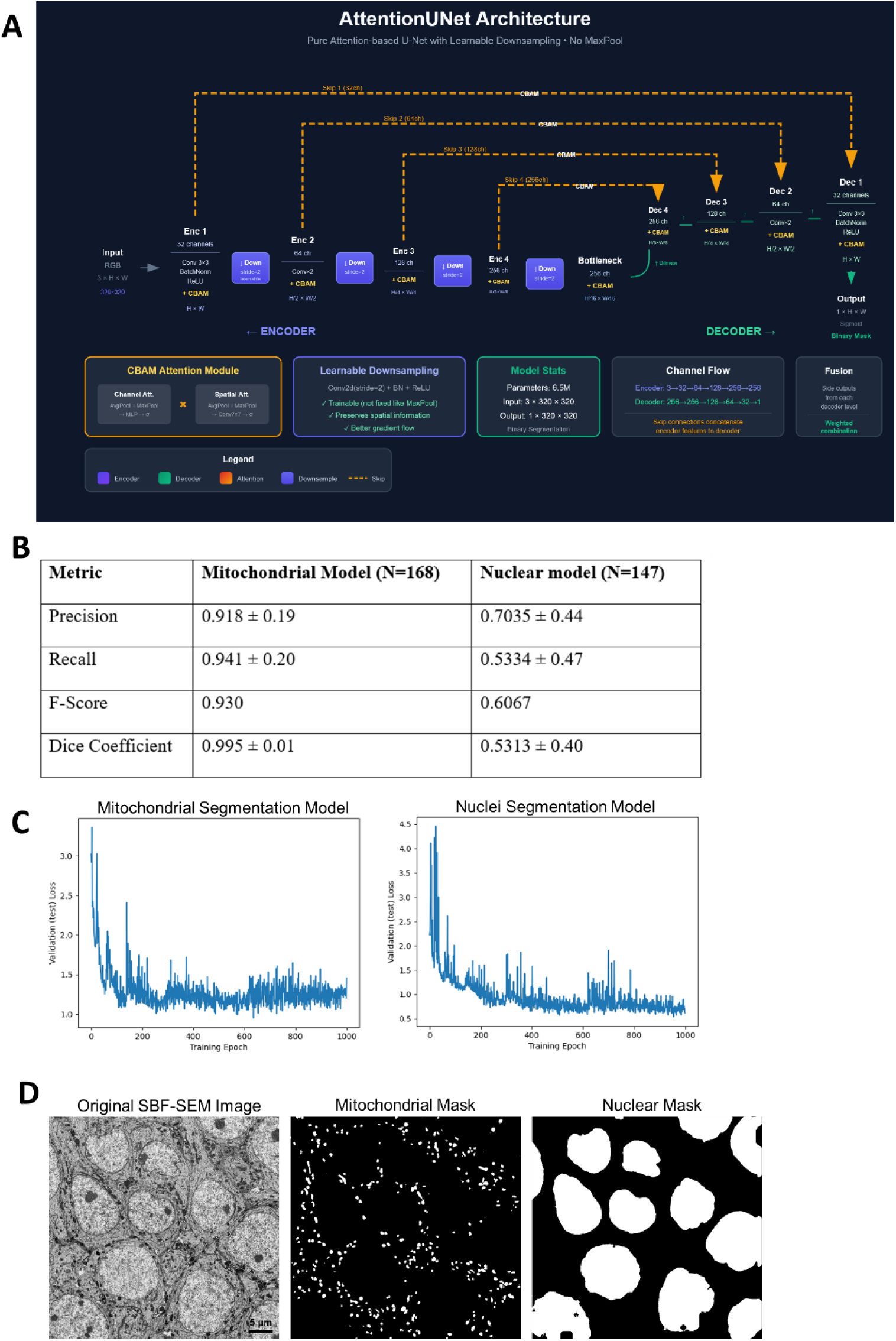
SBF-SEM image segmentation and deep learning–based analysis. (**A**) U-Network Model Architecture: The same model architecture was used for both mitochondrial and nuclear segmentation. (**B**) The table depicts the segmentation performance metrics of the model on the test data for mitochondria (N=168) and nuclei (N=147). Scores are represented as averages with standard deviation based on images that are independent of the training images. Scores were calculated using the Python notebook Metrics.ipnb, which can be found in https://github.com/vipendrakumarsingh/SBF-SEM-image-segmentation. (**C**) Training Loss Curves: The validation loss from the test images during training for the mitochondrial and nuclei segmentation models. (**D**) Binary Masks: An example SBF-SEM image is shown alongside the binary masks generated for the mitochondria and nuclei.

**Figure S2.**
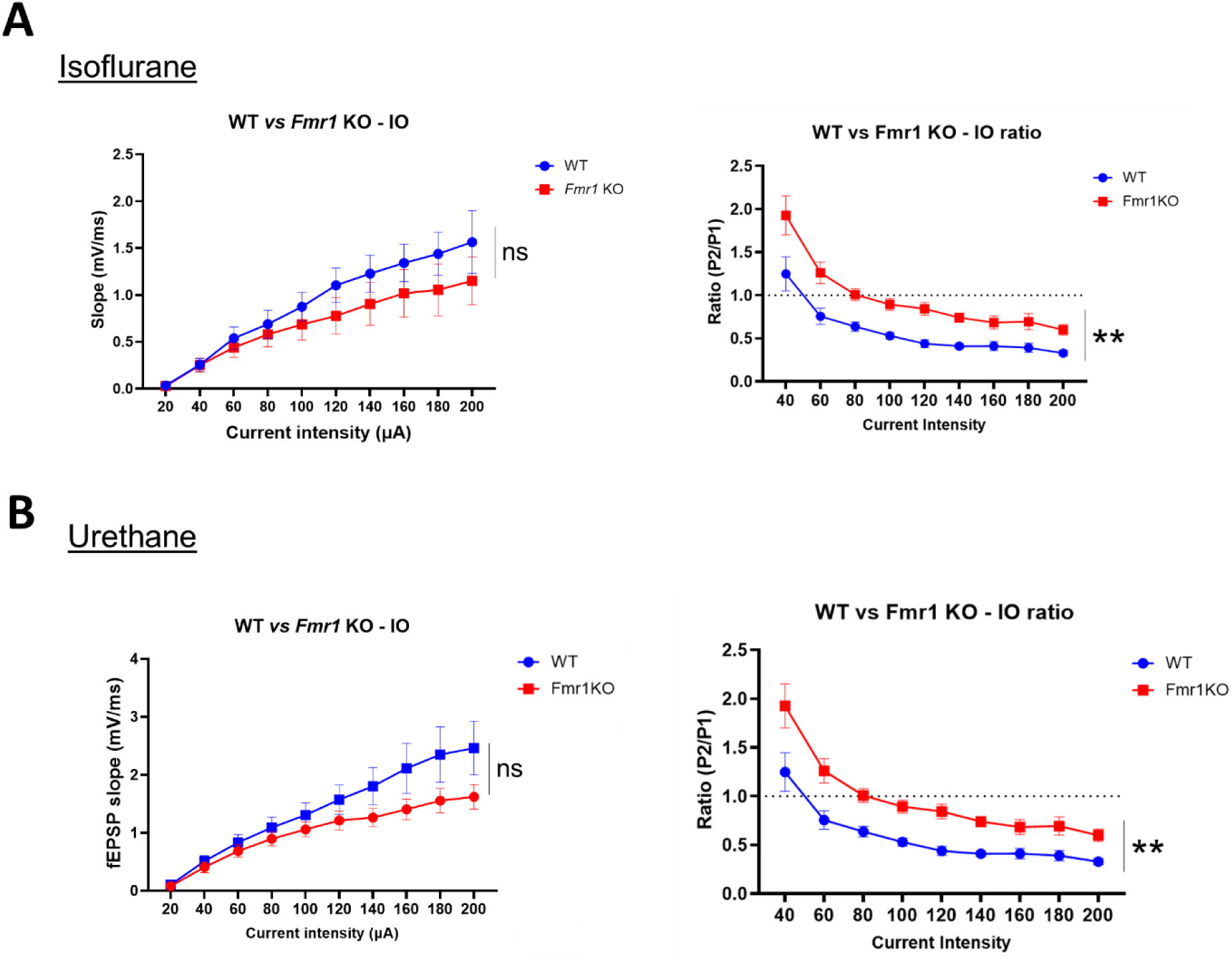
***Fmr1* KO mice exhibit altered paired pulse relations in hippocampal CA1 subfield of isoflurane and urethane anesthetized mice.** (**A**) Input–output curves of fEPSP slope and paired pulse ratios of evoked potentials (EPs) (inter-stimulation interval: 20 ms) from isoflurane anesthetized WT (n = 5) and *Fmr1* KO (n = 6) mice. (**B**) Input–output curves of fEPSP slope and paired pulse ratios (ISI: 20 ms) of EPs from urethane (1.2 g/kg, i.p.) anesthetized WT (n = 10) and *Fmr1* KO (n = 10) mice. No difference in basal synaptic neurotransmission was found between the genotypes when recording EPs in either isoflurane or urethane anesthetized mice. However, *Fmr1* KO mice exhibited a significant increase in paired pulse ratios when compared to WT mice for all tested stimulation intensities in both isoflurane and urethane anesthetized mice. Two-way ANOVA was used to calculate statistical significance. Data are presented as mean ± SEM; ** *p* < 0.01, ns: non-significant.

**Movie S1:** Animation video showing the 3D reconstruction of the cell body of a WT CA1 pyramidal neuron along the z-axis.

